# Using a Riemannian elastic metric for statistical analysis of tumor cell shape heterogeneity

**DOI:** 10.1101/2023.06.11.544518

**Authors:** Wanxin Li, Ashok Prasad, Nina Miolane, Khanh Dao Duc

## Abstract

We examine how a specific instance of the elastic metric, the Square Root Velocity (SRV) metric, can be used to study and compare cellular morphologies from the contours they form on planar surfaces. We process a dataset of images from osteocarcoma (bone cancer) cells that includes different treatments known to affect the cell morphology, and perform a comparative statistical analysis between the linear and SRV metrics. Our study indicates superior performance of the SRV at capturing the cell shape heterogeneity, with a better separation between different cell groups when comparing their distance to their mean shape, as well as a better low dimensional representation when comparing stress statistics. Therefore, our study suggests the use of a Riemannian metric, such as the SRV as a potential tool to enhance morphological discrimination for large datasets of cancer cell images.

## 1 Introduction

Cells cultured on planar surfaces adopt a variety of morphological shapes, that are tightly coupled with molecular processes acting on the cellular membrane and cytoskeleton [19]. This tight coupling suggests various potential applications of quantitative measurements of cellular morphology, e.g. in morphogenesis [26], morphological screening [12] and image guided medical diagnosis, for example to improve the accuracy of cancer tumor grading by cytologists [22]. Using textural or boundary image information, a vast number of features can possibly be used in this context, ranging from human-crafted features to basis function expansions [2,21]. Interestingly, biophysical measurements of cell elasticity in cancer cell have also shown some heterogeneity across cell types, as well as cultures and measurement conditions, which makes their use for diagnosis and treatment promising, yet challenging [9].

In this paper, we consider the so-called *elastic metric*, a Riemannian metric that aims to quantify local deformations of curves by evaluating how they bend and stretch. As a natural geometric tool to investigate the elasticity of cell shape, we apply it to a dataset of tumor cell images that include different cell lines and conditions. Upon processing this dataset to extract cell shapes and using a specific instance of the elastic metric, namely the square root velocity metric (SRV), we find that in comparison with the linear metric, the SRV is superior at capturing and representing the cell shape heterogeneity. Therefore, our study suggests that the elastic metric can be a potential tool to enhance morphological discrimination for heterogeneous datasets of cancer cell images.

## 2 Background

### 2.1 Elastic metric and Square Root Velocity metric for planar curve comparison

#### Definition

The family of *elastic metrics*, introduced by Mio *et al*. [13], can be defined over the space 𝒞 of smooth parametrized curves *c* : [0, 1] *1→* ℝ^2^ with nowhere-vanishing derivative. With *a, b >* 0 denoting the parameters of the family, one associates with every curve *c ∈* 𝒞 an inner product 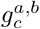 over the tangent space *T*_*c*_𝒞, given by [3,18]

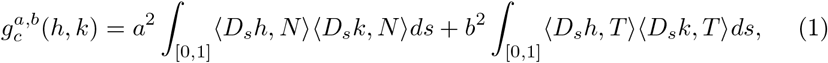

where *h, k* are two curve deformations in the tangent space *T*_*c*_𝒞, that can also be considered as planar curves [13]; *<, >* is the Euclidean inner-product in ℝ^2^, 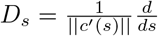, is a differential operator with respect to the arc length *s*, and *N* and *T* respectively are the local unit normal and tangent from a moving frame associated with *c*. Intuitively, elements in *T*_*c*_𝒞 represent infinitesimal deformations of *c*, and 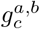 quantifies the magnitude of these deformations, with the two factors *a* and *b* that can be interpreted as weights penalizing the cost of bending (for *a*) and stretching (for *b*) the curve *c*. In this paper, we specifically consider the case (*a, b*) = (1, 1*/*2) that defines the so-called *Square Root Velocity metric*, as it allows in practice for an efficient evaluation [23,10]. In Figure 1.A, we illustrate how the metric can be interpreted for a local deformation *h* of *c*: As we project the derivative of *h* (with respect to its arc length) along the tangent and normal vectors of the reference frame associated with *c*, increasing the bending in *h* results in a relatively higher contribution from the normal component, and thus the integral weighted by *a*^2^, according to Eq. (1). Similarly, stretching increases the contribution from the tangent component, and the integral weighted by *b*^2^.

**Figure 1:**
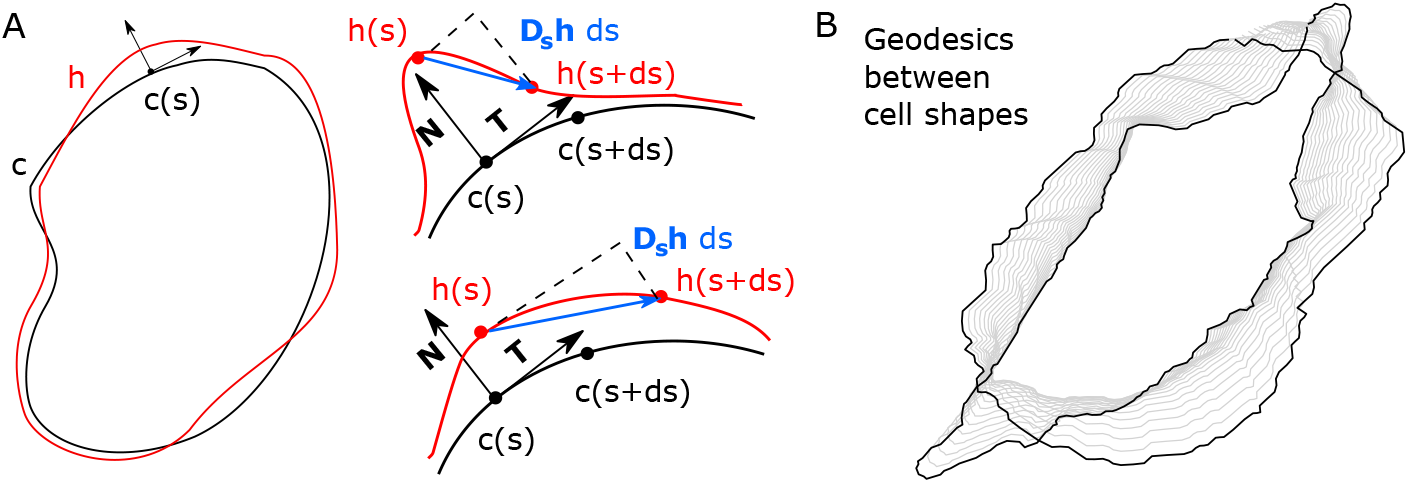
Elastic metric on cell shapes: (**A**) We illustrate how the elastic metric applies to a given shape *c* (shown in left) and a local deformation *h*. According to Eq. (1), this metric is given by the sum of two components, which integrate the projection of the derivative of *h* with respect to the arc length (**D**_**s**_**h** *ds*), on **N** and **T** respectively, which are the local normal and tangent vectors of *c* (shown in right). The projection on **N** (**T**) emphasizes bending (stretching) deformations, as shown in top (bottom) right. (**B**) Upon implementing the metric in Geomstats, we can construct a geodesic path between two cell shapes, as a continuous deformation (with intermediate cells in grey) that minimizes the path length (see Eq. (3)) and yields a geodesic distance (see Material and Methods).

*Geodesic distance:* As a Riemaniann metric [13,23], the elastic metric yields a geodesic distance over 𝒞: For two curves *c*_0_ and *c*_1_ and a regular parameterized path *α* : [0, 1] *1→* 𝒞 such that *α*(0) = *c*_0_ and *α*(1) = *c*_1_, the length of *α*, associated with the elastic metric *g*^*a,b*^ is given by

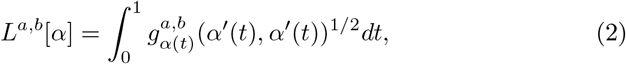

and the *geodesic distance* between *c*_0_ and *c*_1_ is

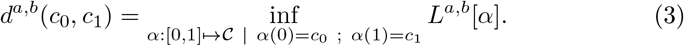

Figure 1.B illustrates the shortest path joining two cell shapes using the elastic metric.

*Fréchet mean:* With the space of curves equipped with this distance, the so-called Fréchet mean of *n* curves (*c*_1_, …, *c*_*n*_) [16] is defined as

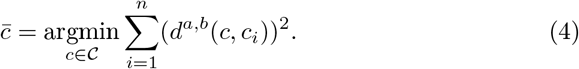

Note that the existence and uniqueness of the Fréchet mean is a priori not guaranteed, but requires that the data is sufficiently concentrated (which we will assume for our datasets) [8].

*Comparison with Euclidean linear metric:* We compare the performance of the elastic metric on cell shapes with the Euclidean linear metric, associated with the ℒ_2_ distance in ℝ^2^. For two cell contours *c*_0_ and *c*_1_ defined as above, this linear distance is given by 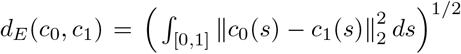, and the *linear* mean of *n* curves 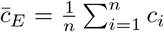 In practice, we compare the use of both metrics in the space of curves quotiented by the action of the group of rigid transformations (i.e. scaling, translation, rotation), via optimal alignment of curve elements [10], using the recent implementation from the *Geomstats* Python library [14] (see Section 3.1).

*Implementation:* An approximation of the geodesic distance associated with the elastic metric *g*^*a,b*^ can be computed as a pull-back of the linear metric: Upon applying a transformation that maps the geodesic associated with *g*^*a,b*^ into a straight line, the geodesic distance is equal to the ℒ^2^ distance between the two transformed curves [18]. While the procedure to construct the mapping can be numerically unstable [3,18], it is simple for the SRV, with the geodesic distance being the ℒ^2^ distance obtained upon representing the curve by its speed, renormalized by the square root of its norm as 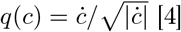

### 2.2 Related works

The elastic metric has been the object of several theoretical and computational developments that primarily focused on curves (for a recent overview, see [4]), with applications to various kinds of biological shapes, including tumor images from MRI (but no cell shapes), plant leafs, or protein backbones [5,6,23]. More recent studies have applied the elastic metric to cell shapes [15,7,17,11] for classification [15,7], dimensionality reduction [11] and regression with metric learning [17]. To our knowledge, the present study is the first to perform a comparative statistical analysis of the elastic metric for tumor cells across different conditions. More generally, there has been a vast number of approaches and features that have been used to model and study cell and/or nuclear shape, as reviewed in [21]; however none of them uses the elastic metric. Overall, the interdependence and relative complexity of the parameters that are classically used to describe biological cells [1,19] make the search of simple interpretable features, such as the elastic metric, useful. For example, a model with two independent nondimensional parameters, that respectively capture flatness and scale, was recently proposed to analyze nucleus shapes of populations for multiple cell lines [2].

## 3 Methods

### 3.1 Datasets

Our dataset consists of images of two murine osteosarcoma cell lines, DUNN and DLM8, which were also previously used for data analysis of cell images [1]. These two cell lines are closely related except for their degree of cancer invasiveness, as the DLM8 line is derived from the DUNN cell line with selection for metastasis. We here consider these two lines separately and present the main results of our statistical analysis for the DUNN line, with results for the DLM8 line being shown at [this link](https://github.com/wxli0/dyn/blob/main/dyn/manuscript/GSI2023_SI.pdf). Both lines have been either treated with DMSO (control) or by cytochalasin D (cytD) or jasplakinolide (jasp). cytD and jasp are two cancer drugs that differently affect the cellular morphology, as cytD leads to actin depolymerization, while jasp enhances it [1]. More details about the experimental methods are available in [1]. We remove outliers from artefacts due to bad segmentation [11], and discretize the cell contour into 100 2D points. After processing, the DUNN cell lines contains 392 cells, including 203 cells in the control group, 96 cells treated by jasp and 93 cells treated by cytD. Similarly, the DLM8 cell line contains 258 cells, including 114 cells in the control group, 82 cells treated by cytD and 62 cells treated by jasp. The datasets are publicly available at this link) and have been added to Geomstats [14]. We preprocessed the datasets by scaling, translation and rotation of the curves: For scaling, we simply normalized the length of each curve to one. For translation, we centered the curves around the plane’s origin. For rotation, we aligned every curve to a reference by finding an optimal rotation to minimize the *L*^2^ distance between two curves. Note that the reparameterization is approximately invariant to the starting points as we selected 200 candidate starting points for each cell when computing the optimal alignment.

### 3.2 Experiments

Our study compares the performance of the SRV metric with the linear metric on analyzing the datasets of tumor cell images, illustrated in Figure 2.A). To do so, we evaluated the associated pairwise distances between cells from Eq. (3), as well as their distance to the mean shape from Eq. (4), upon removing the action of translations and rotations by optimally aligning the curve elements for both the linear and SRV metrics. The analysis is done in Python 3.8 and using the implementation of the SRV in Geomstats [14] with the version from Aug 23, 2022, with scripts reproducing our results available in this [github repository](https://github.com/wxli0/dyn).

**Figure 2:**
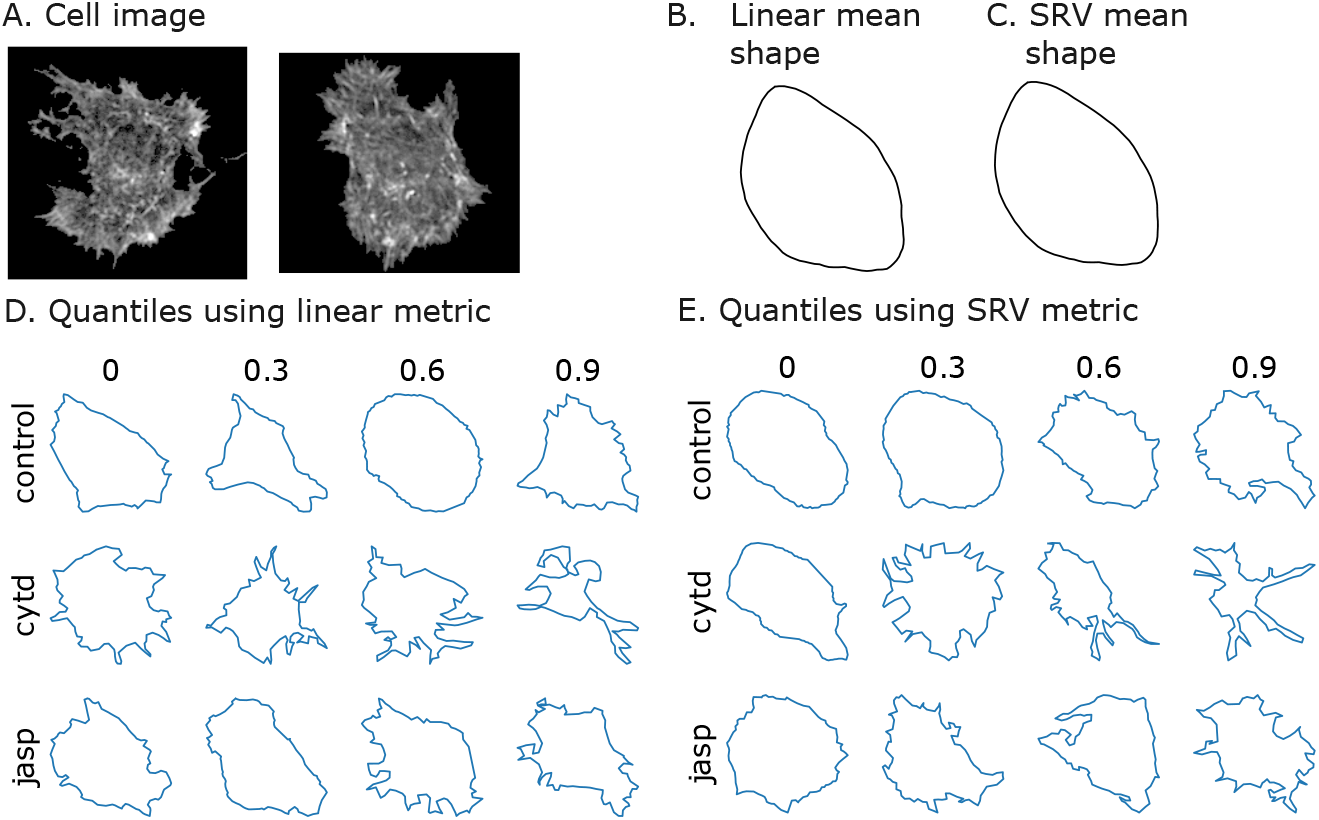
**A:** Two examples of osteocarcoma cell image obtained from fluorescence microscopy. **B:** Mean cell shapes computed over the cells from the DUNN cell line using the linear metric. **C**: Same as **(B)** with the SRV metric. **D:** Quantiles of distance to the mean shape for different conditions using the linear metric (dataset of DUNN cells). **E:** Same as (D) for the SRV metric.

## 4 Results

### 4.1 Visualization of shape quantiles

Upon considering the linear and SRV metrics, we evaluate for each cell of the dataset the distance to the mean shape associated with each metric (the mean shapes can be visualized in Figures 2.B and C. For the three conditions that reflect different treatments applied to the cells (see section 3.1), we visualize in Figures 2.D and E the quantiles associated with the 0th, 30th, 60th and 90th percentiles of the resulting distance distribution. The results suggest that while the linear and SRV mean shapes are quite similar, the SRV metric captures irregularities in shape, as increasingly irregular shapes appear as the quantile increases in Figure 2.E, with farthest cells having large invaginations and overhangs in the cell perimeter, which are captured in the “bending” term of the SRV metric (Eq. (1)). In comparison, the linear metric detects less regular cells on the 0th percentile for cells in all control and treatment groups. We also obtained similar findings with the DLM8 cell line (data not shown here, available at [this link](https://github.com/wxli0/dyn/blob/main/dyn/manuscript/GSI2023_SI.pdf)).

### 4.2 Histograms of distances to the mean

To confirm the visual impression from Figure 2, we plot in Figure 3 the histograms of distances from the global mean for the different conditions (control and treatments). Upon comparing the histograms produced by the linear (Figure 3.A and SRV (Figure 3.B) metrics (defined in section 2.1), we find that while the linear metric does not indicate significant differences between the jasp and control conditions, the SRV metric globally yields a better separation across the conditions (with similar results for the DLM8 cell line at [this link](https://github.com/wxli0/dyn/blob/main/dyn/manuscript/GSI2023_SI.pdf)). Using the SRV metric, the distribution of cytD cells is shifted to the right, while jasp treated cells are shifted to a less extent, yielding a narrower distribution. Interestingly, these observations are in line with the biological mechanisms of action of cytD and jasp: As cytD disassembles actin filaments and causes cytosolic aggregation of actin masses [25], the resulting loss of internal tension forces is expected to give rise to shapes with many invaginations, and increasing distance from the smooth ellipsoid. In comparison, the effects of jasp should be less dramatic since it leads to the formation of a more diffuse actin network, but without affecting stress fibers [20]. The SRV metric also remarkably yields a bimodal distribution for the control cells, which possibly suggests the existence of multiple states within the cell line. To visualize the difference between these the two modes identified by the SRV metric, we show the mean cell shapes over the subsets of cells in the bins 0.06-0.08 (Figure 3.C) and 0.18-0.20 (Figure 3.D) of the histogram, which correspond to the two observed peaks. The two mean cells are remarkably different, with the edge of mean cell in bin 0.06-0.08 being smooth and the edge of the mean cell in bin 0.18-0.20 being rough. This observation is expected since we previously found that cells that are farther away from the mean are more irregular. Using the DLM8 cell line, we also find a better (but less significant) separation across the conditions with the SRV metric (link [here](https://github.com/wxli0/dyn/blob/main/dyn/manuscript/GSI2023_SI.pdf)).

**Figure 3:**
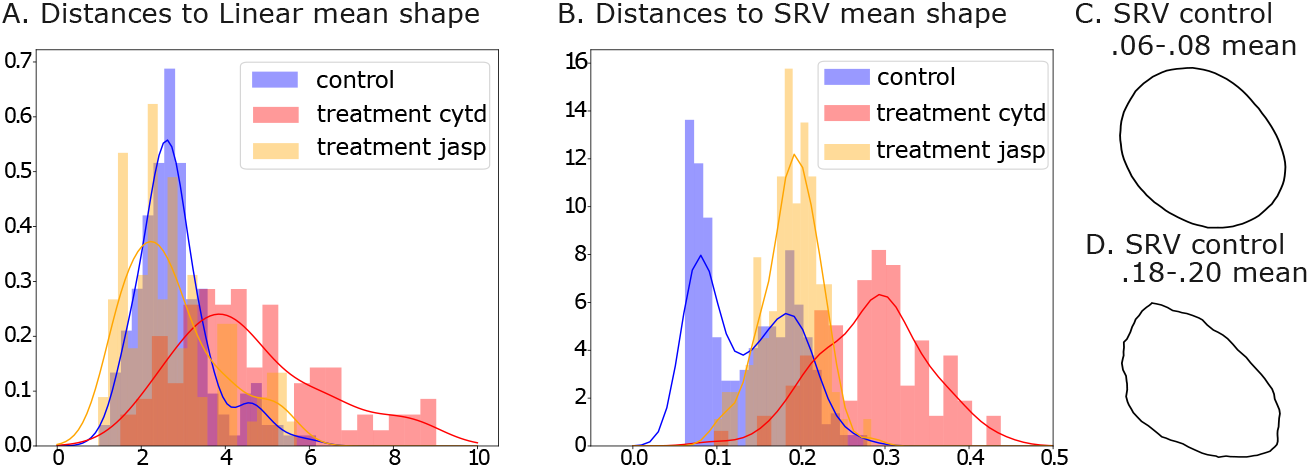
Histograms of distances to linear mean (**A**), and SRV Fréchet mean (**B**) for the DUNN cell line. We observe that the distances of cells in the control group are closer than those in the treatment groups using SRV metric. The curves present kernel-density estimate for each group using Gaussian kernels. Mean cell shapes of control cells with SRV Fréchet mean in 0.06-0.08 (**C**), and with SRV Fréchet mean in 0.18-0.20 (**D**) using the SRV metric.

### 4.3 Visualization in lower dimension space

As dimensionality reduction methods are usually employed to visualize and interpret large datasets of cell images [1], we finally compare the results obtained from projecting the data into a lower dimension space using both metrics. To do so, we perform a multidimensional scaling (MDS) on the pairwise distance matrix, with the results for the DUNN line shown in 2D (Figure 4) and 3D (results available [here](https://github.com/wxli0/dyn/blob/main/dyn/manuscript/GSI2023_SI.pdf)). Our results suggest that the SRV metric tends to better capture the cell shape heterogeneity, as cells are overall more spread in both 2D and 3D, with the control group mostly being centered in the middle, surrounded by jasp treated cells and cytD treated cells located further. We also examine the stress statistics obtained for the different dimensions tested in MDS, as an indicator of the goodness-of-fit [24]. The MDS always achieves a better stress statistic for the SRV and all dimension tested (results available [here](https://github.com/wxli0/dyn/blob/main/dyn/manuscript/GSI2023_SI.pdf)), suggesting that the metric captures more informative patterns when it comes to dimensionality reduction. We obtain similar results for the DLM8 line (link [here](https://github.com/wxli0/dyn/blob/main/dyn/manuscript/GSI2023_SI.pdf)).

**Figure 4:**
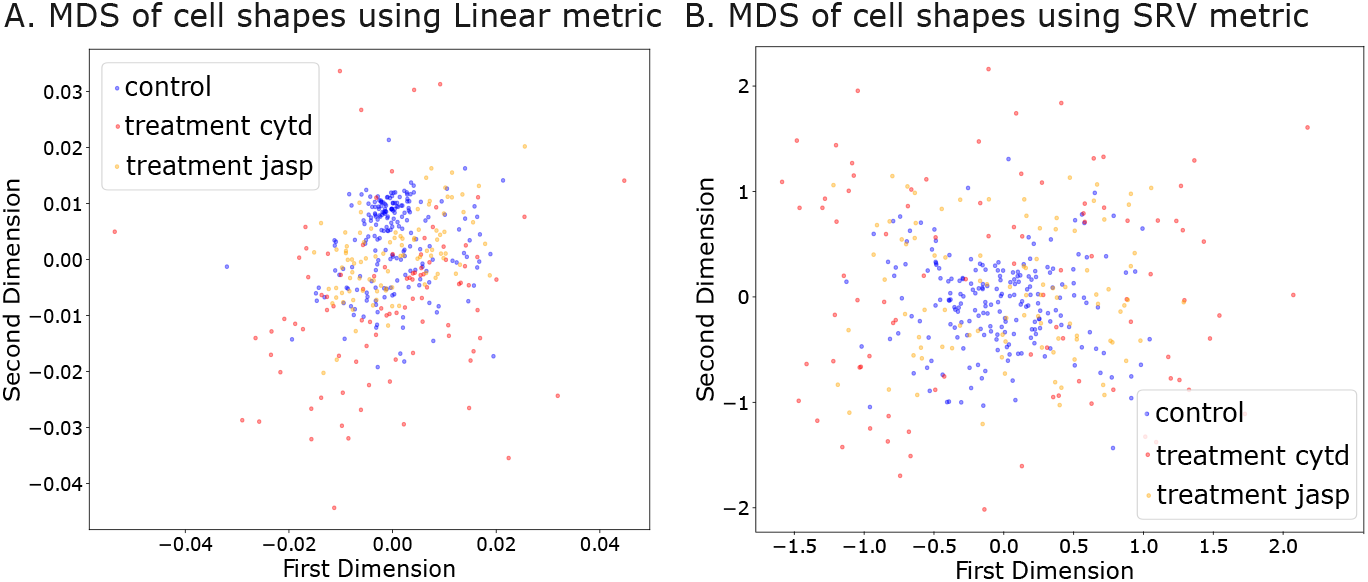
Projections for MDS in 2D using the linear metric **(A)**, and the SRV metric **(B)**. We observe that cells in different control and treatment groups are more uniformly spread using the SRV metric in 2D.

## 5 Conclusion

We used the SRV metric to statistically analyze 2D shapes from tumor cell images, and showed some superior performance over the linear metric in all the tasks tested. The results of our comparative analysis were consistent across different cell lines of our datasets, suggesting that the elastic metric is better suited for interpreting morphological changes in the cell shape. While our study suggests the potential use of this Riemaniann metric for cancer cell detection, it is still limited to a relatively small dataset. The development of the elastic metric as a tool for practitioners requires some more thorough investigation of the morphological changes observed across different cell types and conditions, jointly with other features of the literature. The evaluation of the SRV distance in our paper could also be refined, by notably quotienting out the action of reparametrization [10,4], and it would be interesting to generalize our results to the whole family of elastic metrics, and compare their performance with some alternative metrics. In particular, considering the *H*_1_ metric would be useful to determine if the performance of the elastic metric comes from its non-linear nature, or from the derivative information it contains. We are currently pursuing these directions.

