## Supplemental Information for "Using a Riemannian elastic metric for statistical analysis of tumor cell shape heterogeneity"

This Supplemental Information File contains supplementary figures S1-4 to our main manuscript, showing results of our study on the DLM8 cell line and additional results for the DUNN cell line.

### Supplementary Figures

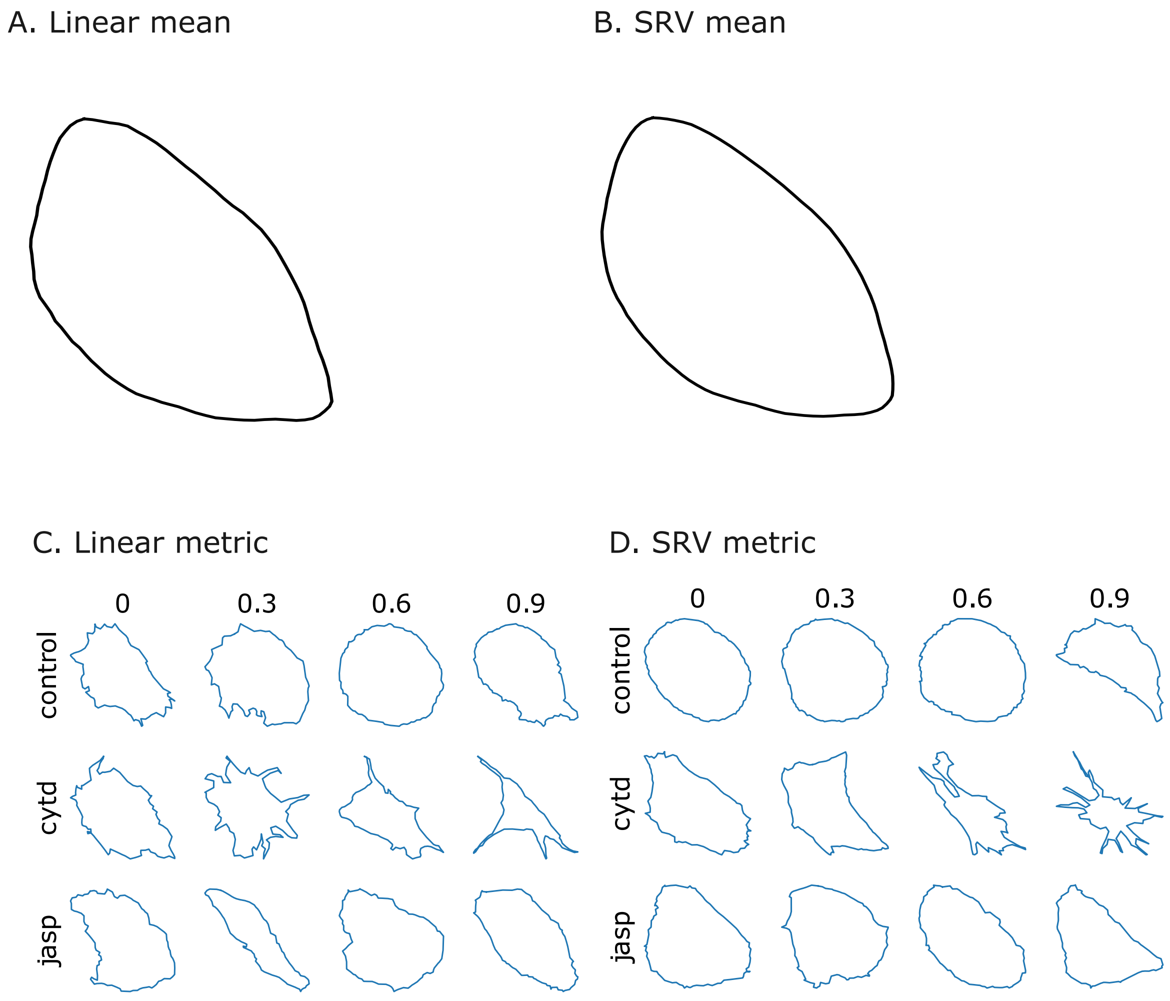


**Figure** S1: Mean shape and quantile visualization for the DLM9 cell line. **A:** Mean cell shape obtained using the linear mean. **B:** Mean cell shape obtained using the SRV mean. **C:** Quantiles of distance to the mean shape for different conditions using the linear metric . **D:** Same quantile visualization as in (C) for the SRV metric. Note that more regular cells are placed on the 0 percentile for cells in all control and treatment groups using the SRV metric.

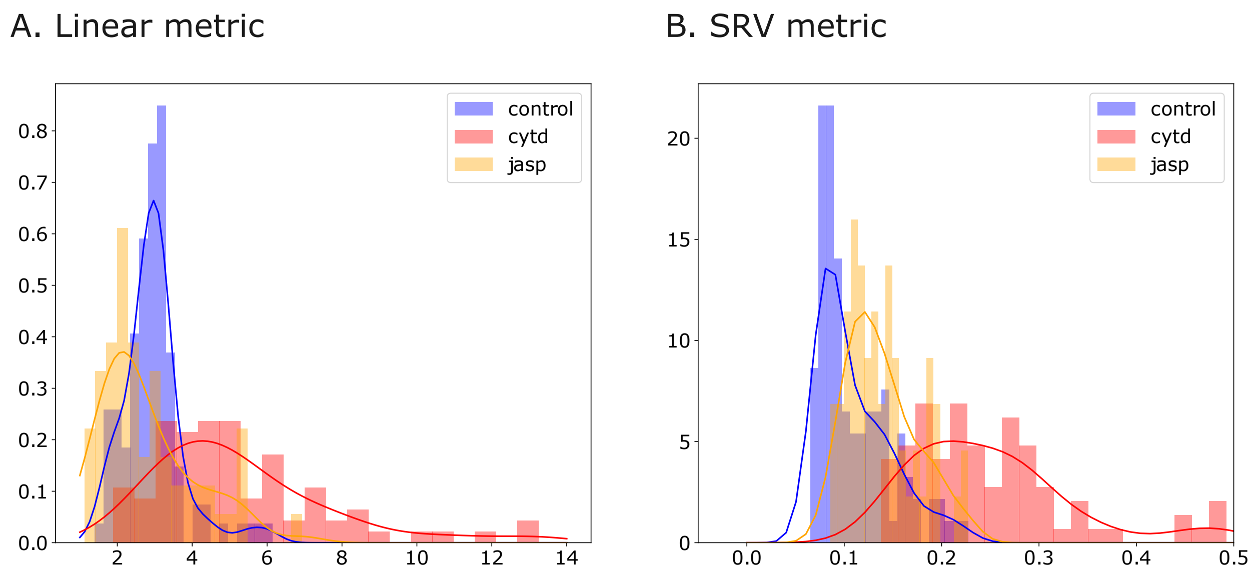


**Figure** S2: Histograms of distances for the DLM8 cell line to the linear mean (**A**), and the SRV mean (**B**). We observed that cells in different control and treatment groups are more uniformly spread using the SRV metric in 2D. We observed that the distances of cells in the control group are closer than those in the treatment groups using SRV metric. The curves present kernel-density estimate for each group using Gaussian kernels.

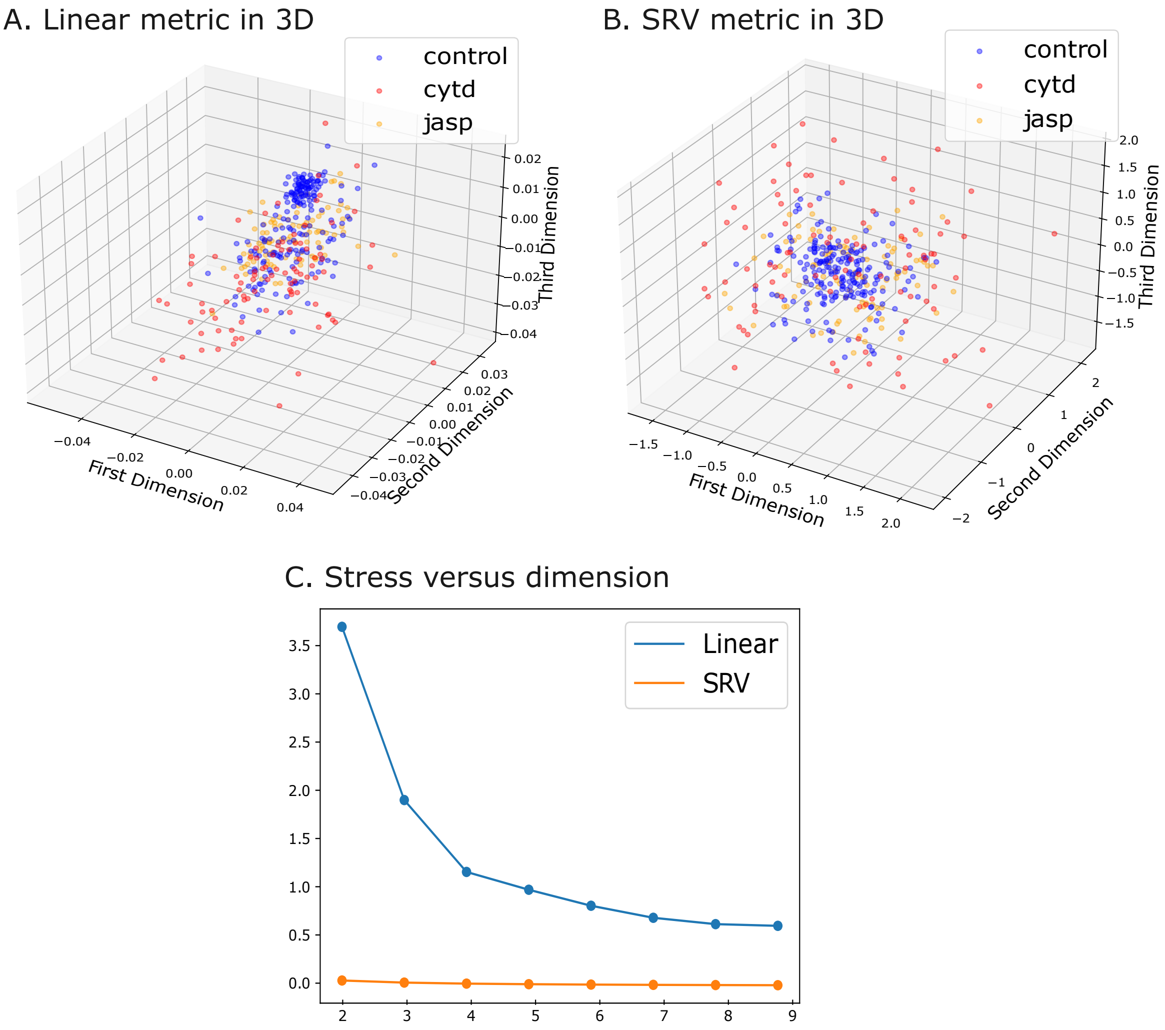


**Figure** S3: MDS of cell shapes and stress statistics from the DUNN cell line. **A:** MDS projection in 3D using the linear metric. **B:** MDS projection in 3D using the SRV metric. We observe that cells in different control and treatment groups are more uniformly spread for the SRV metric in 3D. **C:** Stress versus dimension for DUNN cell lines using the linear metric (blue) and SRV metric (orange). We observed the elastic metric always achieves a lower (better) stress statistic than the linear metric.

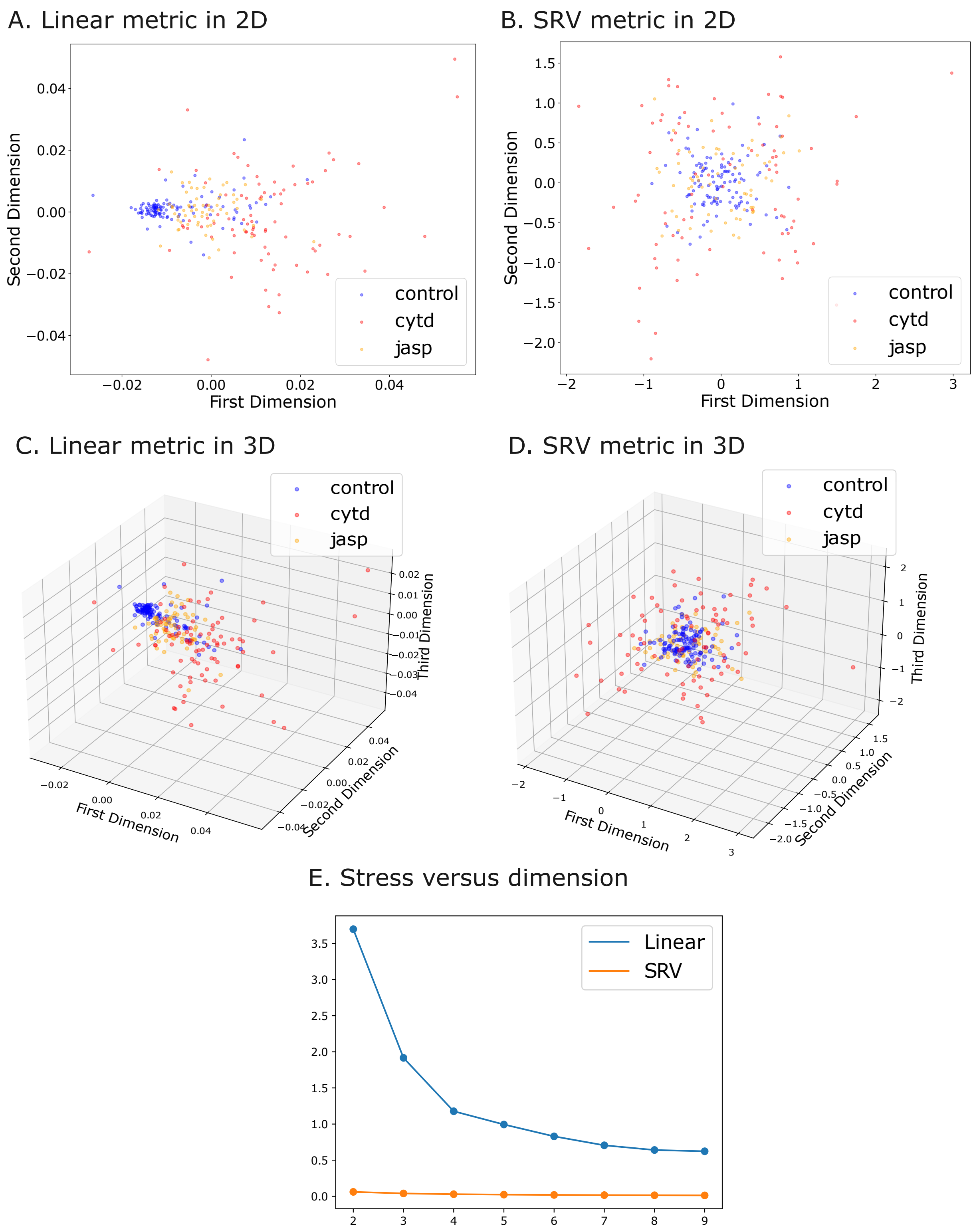


**Figure** S4: MDS of cell shapes and stress statistics from the DLM8 cell line. **A:** MDS projection in 2D using the linear metric. **B:** MDS projection in 2D using the SRV metric. **C:** MDS projection in 3D using the linear metric. **D:** MDS projection in 3D using the SRV metric. **E:** Stress versus dimension for DLM8 cell lines using the linear metric (blue) and the SRV metric (orange). We observed the elastic metric always achieves a lower (better) stress statistic than the linear metric.
